# A chemogenetic CRISPR knockout screen identifies a ubiquitin-mediated calcineurin-BMP pathway relay in T cell activation

**DOI:** 10.64898/2026.07.08.737199

**Authors:** Marcel Diallo, Leon Müller, Stefanie Uhlig, Ivo Hendriks, Julia Kzhyshkowska, Harald Klüter, Michael Nielsen, Jesper Olsen, Patrick Wuchter, Karen Bieback, Jakob Nilsson, Gianmatteo Vit

## Abstract

T cell activation is dependent on calcineurin signalling, yet how this pathway integrates with other regulatory systems remains incompletely understood. Here, we exploit the dual pharmacology of FK506, which inhibits calcineurin while also releasing FKBP12-mediated repression of BMP receptors, to dissect signalling crosstalk during T cell responses. A comparative genome-wide CRISPR knockout screen using FK506 and Cyclosporin A revealed that FK506 uniquely engages BMP pathway components and ubiquitin regulatory networks. Functional analyses in Jurkat and primary human T cells showed that FK506 induces BMP receptor-dependent activation of SMAD1/5/8 and triggers a rapid remodelling of K48- and K29-linked ubiquitin chains. Proteomics further demonstrated selective ubiquitination of immune signalling proteins and ubiquitin regulators, linking BMP pathway activation to proteasome-dependent turnover and non-proteolytic ubiquitin signalling. In primary T cells, BMP receptor signalling enhance calcineurin-driven activation, while changes in ubiquitin conjugation modulate this response, thereby establishing an integrated calcineurin-BMP-ubiquitin relay. Together, our results uncover a context-dependent signalling axis in which BMP activation and ubiquitin dynamics fine-tune calcineurin-mediated T cell signalling.

## INTRODUCTION

T cell activation is governed by the integration of multiple signalling pathways that coordinate transcriptional programs, metabolic reprogramming, and effector functions. Among these pathways, calcium-dependent activation of the phosphatase calcineurin and subsequent nuclear translocation of NFAT (Nuclear Factor of activated T cells) transcription factors represents a central axis controlling T cell-driven immune responses (Clipstone and Crabtree, 1992; Trebak and Kinet, 2019). Pharmacological inhibition of calcineurin using the immunosuppressive drugs tacrolimus (FK506) and Cyclosporin A (CsA) remains a cornerstone of clinical immunosuppression in transplantation and autoimmune diseases (Halloran, 2004).

Although FK506 and CsA both suppress T cell activation through calcineurin inhibition, they use distinct molecular mechanisms. CsA forms a ternary complex with cyclophilin A and calcineurin, whereas FK506 binds FKBP12 (FK506-Binding Protein 12), and the resulting complex inhibits calcineurin phosphatase activity (Kissinger *et al*., 1995; Ho *et al*., 1996). Beyond calcineurin inhibition, FK506 possesses additional molecular activities that are absent in CsA. Importantly, FKBP12 acts as a negative regulator of bone morphogenetic protein (BMP) signalling by binding to the glycine-serine region of the BMP type I receptor, thereby preventing its activation. FK506 binding to FKBP12 releases this inhibitory constraint, resulting in potentiation of BMP signalling through enhanced SMAD1/5/8 phosphorylation (Huse *et al*., 1999; Spiekerkoetter *et al*., 2013).

BMP signalling, a branch of the transforming growth factor-β (TGF-β) superfamily, has been extensively studied in development, tissue homeostasis, and disease (Richardson *et al*., 2023). While the BMP pathway is increasingly recognized for its roles in non-developmental contexts, including immune regulation, its function in T cell biology remains incompletely understood (Saadey *et al*., 2023). Previous studies have suggested context-dependent roles for BMP signalling in immune cell differentiation and function, but the mechanisms integrating the BMP pathway with canonical T cell activation pathways have remained elusive (Martínez *et al*., 2015). Notably, emerging evidence indicates that calcineurin can directly intersect with the BMP pathway by modulating SMAD1/5 phosphorylation and by regulating the expression of BMP ligands such as BMP2, pointing to a context-dependent interplay between these signalling axes (Cho *et al*., 2014; Hendrikx *et al*., 2019).

Recent work has demonstrated that FK506-mediated BMP pathway activation is independent of calcineurin inhibition and can be preserved in calcineurin-sparing FK506 analogues (Peiffer *et al*., 2019; Larraufie *et al*., 2021). These findings suggest the possibility that FK506 exerts dual and potentially opposing effects on T cells: while calcineurin inhibition suppresses T cell functions, BMP activation may modulate immune processes in a distinct manner. However, whether and how BMP-mediated responses intersect with calcineurin-dependent T cell activation remains unclear.

To address this, we exploited the unique pharmacological property of FK506 as a tool to interrogate the molecular interplay between calcineurin and BMP pathways and performed a genome-wide CRISPR knockout screen in Jurkat T cells using FK506 and, as a control, Cyclosporin A (McKinley and Cheeseman, 2017; Bock *et al*., 2022). By directly comparing these two calcineurin inhibitors, one that activates BMP signalling and one that does not, we aimed to distinguish calcineurin-selective effects from BMP-specific mechanisms. This approach allowed us to identify genetic dependencies associated with BMP pathway engagement and to define how these signals overlap with calcineurin signalling during T cell activation.

Using biochemical, functional, and proteomic approaches in Jurkat cells and primary human T cells, we uncover a calcineurin-BMP signalling relay that is engaged only upon T cell activation and is coupled to extensive remodelling of the ubiquitinome. Our findings establish ubiquitin-dependent BMP signalling as a positive regulator of T cell function and provide a basis for understanding BMP pleiotropic effects upon calcineurin activation in T cell responses.

## RESULTS

### A CRISPR knockout screen with FK506 identifies BMP pathway components and ubiquitination networks as determinants of FK506 response

Given the ability of FK506 to inhibit calcineurin while simultaneously activating BMP signalling, we aimed to identify genetic signatures that reveal functional crosstalk between calcineurin and BMP signalling in regulating T cell activation. To this end, we performed a comparative genome-wide CRISPR knockout screen in Jurkat cells using FK506 and CsA, a mechanistically related calcineurin inhibitor that does not activate BMP signalling, thereby serving as a control. Jurkat E6.1 cells were transduced with the TKOv3 “all-in-one” CRISPR library targeting 18,053 protein-coding genes and, after puromycin selection, were expanded for 6 days before treatment with sublethal doses (LD20) of FK506 or CsA. After 12 days of drug exposure, sgRNA representation was quantified by next-generation sequencing and a normalized depletion score (Z-score) was calculated for each gene, reflecting its impact on cellular fitness under each treatment condition (Figure 1A and S1A-B). The two screens produced markedly different chemogenetic profiles (Figure 1B-C). Cells challenged by FK506 treatment revealed a distinct dependency pattern dominated by BMP pathway components and ubiquitin regulators. Knockout of BMP receptors ACVR2B, BMPR1A, ACVR1, and BMPR2, followed by SMAD4 and its positive regulator AMBRA1, as well as ID3, the downstream target of BMP signalling, produced the strongest resistance and thus conferred a selective advantage to the cells (O’Shaughnessy *et al*., 2004; Schmierer and Hill, 2007; Liu *et al*., 2021). Conversely, disruption of negative regulators of BMP signalling (SMAD6) sensitized cells to FK506, further supporting the notion that FK506 engages BMP signalling to exert its function (Spiekerkoetter *et al*., 2013; Peiffer *et al*., 2019; Larraufie *et al*., 2021). Further analysis of the FK506 chemogenetic profile revealed a prominent enrichment of ubiquitin-related genes among both resistance and sensitivity hits, as observed for BMP regulators. Loss of the HECT-type E3 ubiquitin ligases UBR5 and TRIP12, as well as the E3 ligase adaptor AMBRA1, and the deubiquitinase (DUB) OTUD5 conferred resistance to FK506, whereas knockout of the E3 ligase HECTD1, the E2 ligase SMURF2, the ubiquitin-conjugating enzyme UBE2K and the deubiquitinase ZRANB/TRABID, a HECTD1 stabilizer, resulted in strong synthetic lethality (Kushioka *et al*., 2020; Chaikovsky *et al*., 2021; Harris *et al*., 2021; Kaiho-Soma *et al*., 2021; Naksone *et al*., 2022; Wang *et al*., 2023; Chen *et al*., 2025; Morita *et al*., 2025; Sinha *et al*., 2026). This bidirectional representation of ubiquitin regulators and BMP signalling components suggests that both processes play a central role in shaping the cellular response to FK506 (Figure 1B). The CsA screen did not reveal components of the BMP pathway or ubiquitin-related genes, but instead showed enrichment for unfolded protein response and ER/Golgi-related components, highlighting a gene-dependency profile distinct from that of FK506 (Figure 1C). Together, these data show that FK506 and CsA impose fundamentally different genetic dependencies in T cells, with FK506 uniquely coupling BMP signalling to ubiquitin-dependent regulation.

**Figure 1.**
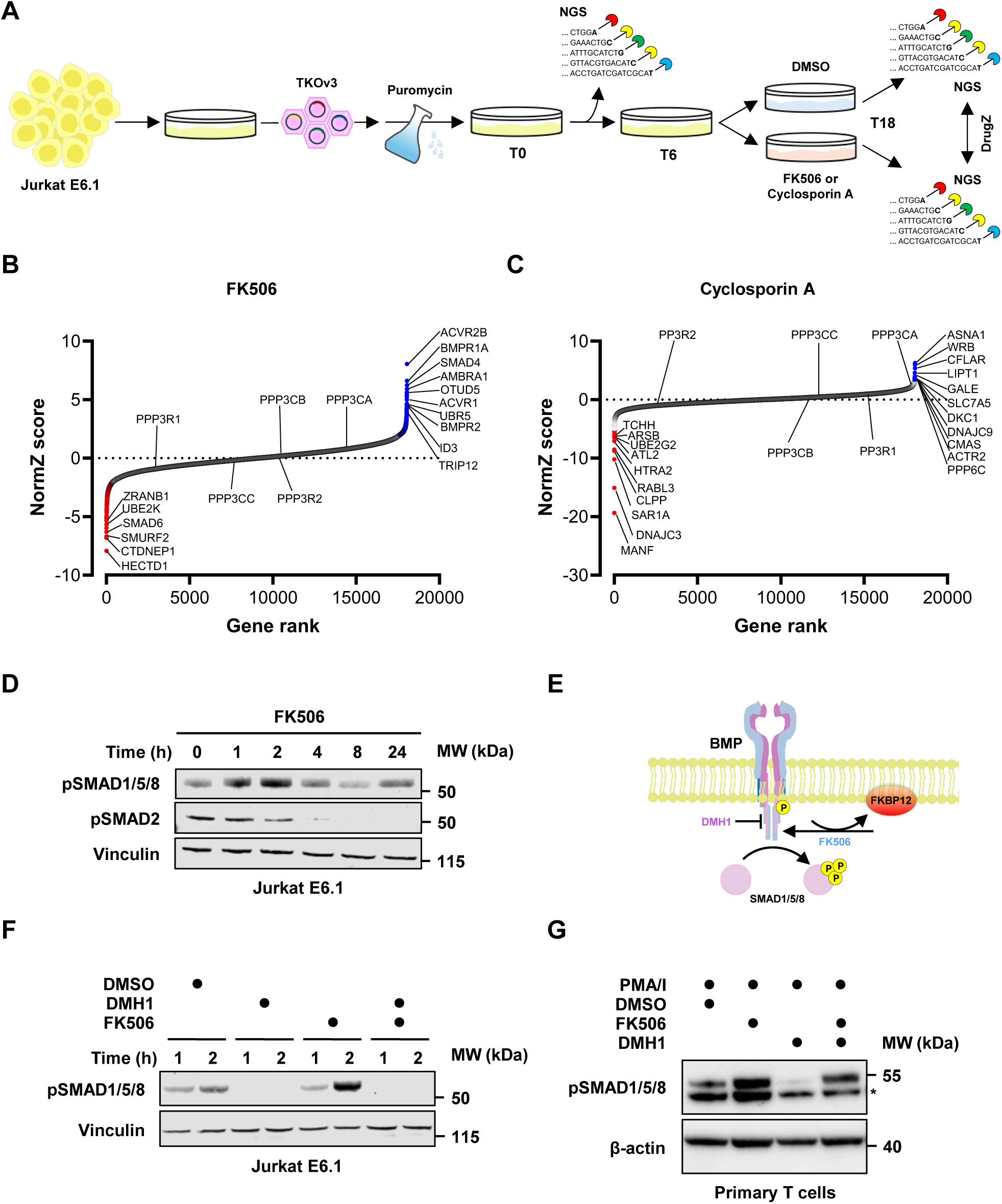
CRISPR screen with FK506 identifies BMP pathway components and ubiquitination networks as determinants of FK506 response. **(A)** Schematic of the genome-wide CRISPR knockout screen. Jurkat E6.1 cells transduced with the TKOv3 sgRNA library were selected with puromycin and sampled at T0 for NGS (baseline). After recovery for 6 days (T6), cells were treated with DMSO, FK506, or Cyclosporin A (CsA) until T18, followed by NGS-based sgRNA quantification and DrugZ software analysis to identify drug response modulators. **(B-C)** Gene ranking based on DrugZ-normalized scores (normZ) from the FK506 and CsA screens. Positive scores indicate resistance (sgRNA enrichment), while negative scores indicate sensitization (sgRNA depletion). Distinct patterns of gene enrichment and depletion are observed after FK506 and CsA treatment. **(D)** Time-course analysis of SMAD1/5/8 and SMAD2 phosphorylation (pSMAD1/5/8 and pSMAD2) in Jurkat E6.1 cells treated with FK506. pSMAD1/5/8 and pSMAD2 were assessed by Western blotting at the indicated time points. **(E)** Schematic representation of BMP receptor signalling. Activation after FKBP12 dissociation from BMP type I receptors upon FK506 treatment leads to SMAD1/5/8 phosphorylation. DMH1 inhibits BMP type I receptor kinase activity. **(F)** pSMAD1/5/8 protein expression in Jurkat E6.1 cells after treatment with FK506 or DMH1, alone or in combination. **(G)** pSMAD1/5/8 protein expression in primary human T cells following treatment with PMA/Ionomycin (PMA/I), FK506, or DMH1, as indicated. *, non-specific signal. **(D and F-G)** Vinculin or β-actin were used as loading control. Blots are representative of three independent experiments.

To functionally validate the suggested BMP pathway activation after FK506 treatment, we monitored the phosphorylation of SMAD1/5/8 (pSMAD1/5/8) and SMAD2 (pSMAD2) as markers of BMP and TGF-β signalling, respectively. In Jurkat cells, FK506 treatment resulted in increased pSMAD1/5/8 within one hour, accompanied by a reduction in pSMAD2 levels (Figure 1D). Importantly, FK506-induced SMAD1/5/8 phosphorylation was abrogated by the BMP type-I receptor inhibitor DMH1 (Alsamarah *et al*., 2015) (Figure 1E-F). We recapitulated this effect in primary human T cells upon activation with PMA and Ionomycin (PMA/I), which activate the Ca²⁺/calcineurin axis (Chatila *et al*., 1989), indicating that FK506-mediated activation of BMP signalling represents a conserved mechanism in cell lines and human primary T cells (Figure 1G).

### BMP signalling regulates ubiquitin dynamics in activated T cells

While the CRISPR screen identified ubiquitin-related genes as key modulators of FK506 response and biochemical assays confirmed FK506-mediated activation of BMP signalling, it was unclear whether these observations converge at the level of ubiquitin-dependent regulation. Given the important role of ubiquitination and BMP pathway in immune regulation, we next sought to determine whether FK506 treatment directly impacts the cellular ubiquitination landscape and whether this effect is linked to BMP receptor activity. To explore this, Jurkat cells were treated with FK506 for up to 24 hours to assess polyubiquitin levels by immunoblotting. FK506 treatment induced a rapid accumulation of K48-linked polyubiquitin chains, peaking at approximately two hours after treatment and gradually declining thereafter. Notably, pSMAD1/5/8 displayed similar kinetics, suggesting a temporal link between BMP pathway activation and remodelling of the ubiquitinome (Figure 2A). To further examine the relationship between BMP signalling and ubiquitination, we tested whether activation of the pathway by its endogenous ligand similarly affects ubiquitin dynamics. Treatment with recombinant BMP7 led to a robust increase in K48-linked polyubiquitin chains, indicating that ubiquitination is a component of BMP pathway activation (Figure 2B). Co-treatment with the BMP type I-receptor inhibitor DMH1 markedly attenuated FK506-induced K48 polyubiquitin accumulation, indicating that FK506-driven ubiquitination is, at least in part, dependent on BMP receptor signalling (Figure 2C). We recapitulated this effect in primary human T cells isolated from healthy donors. Importantly, FK506-induced accumulation of polyubiquitin chains in primary T cells was restricted to PMA/I-activated T cells, whereas no increase in ubiquitination was detected in unstimulated naïve T cells (Figure 2D). In contrast, Jurkat cells displayed FK506-induced ubiquitination in the absence of exogenous stimulation (Figure 2A and 2B), consistent with their intrinsically activated state and limited capacity for further activation by PMA/I (Figure S2A). These findings suggest that BMP-dependent ubiquitin remodelling is largely restricted to activated T cells.

**Figure 2.**
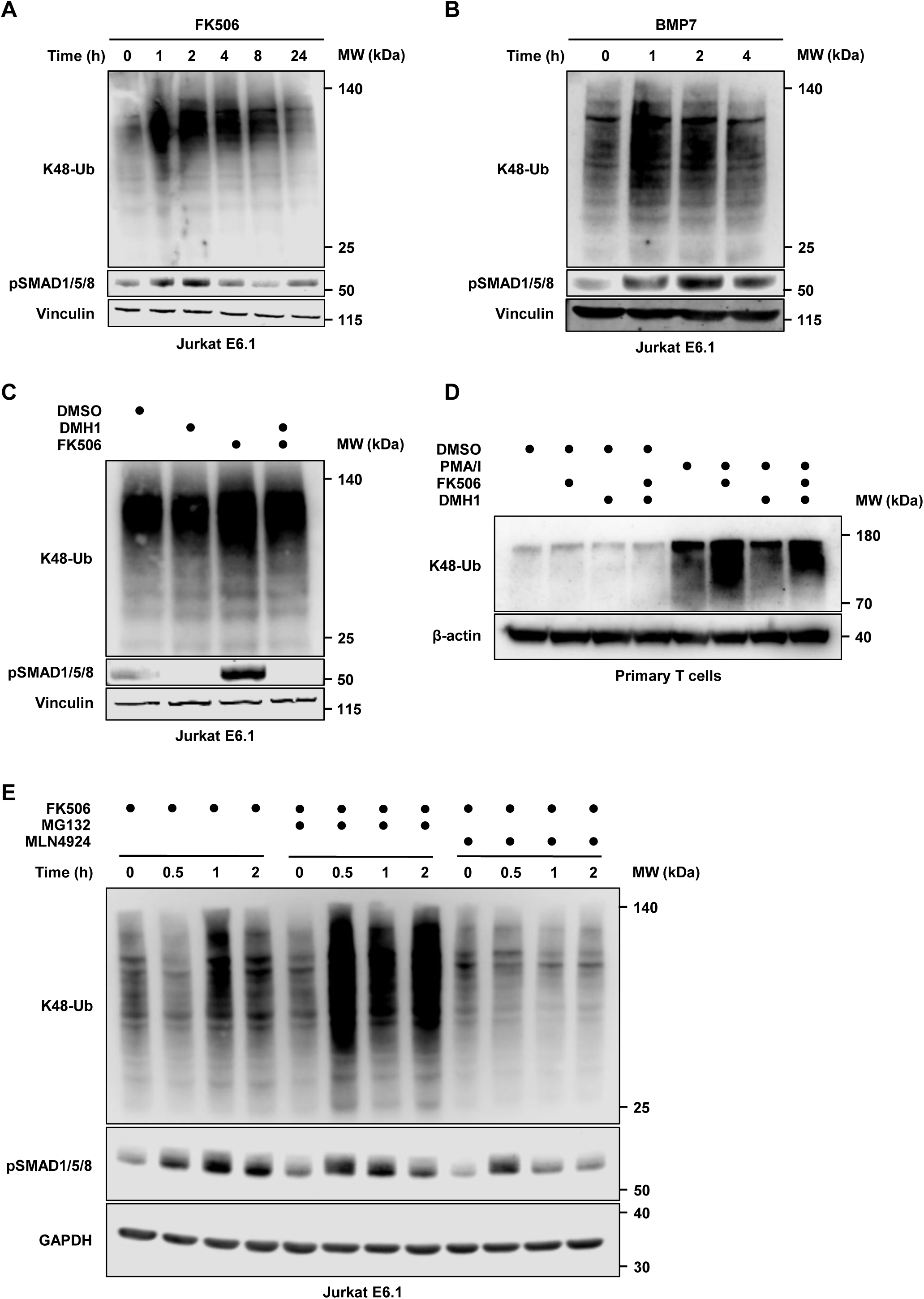
BMP signalling regulates ubiquitin dynamics in activated T cells. **(A)** Time-course analysis of K48-linked polyubiquitin chains (K48-Ub) and pSMAD1/5/8 in Jurkat E6.1 cells treated with FK506 at the indicated time points. **(B)** Time-course analysis of K48-Ub and pSMAD1/5/8 in Jurkat E6.1 cells stimulated with recombinant BMP7 protein. **(C)** K48-Ub and pSMAD1/5/8 levels in Jurkat E6.1 cells treated with FK506 or DMH1, alone or in combination, as indicated. **(D)** K48-Ub levels in primary human T cells treated with PMA/Ionomycin (PMA/I), FK506, or DMH1, alone or in combination, as indicated. **(E)** Time-course analysis of K48-Ub and pSMAD1/5/8 in Jurkat E6.1 cells treated with FK506 alone or in combination with the proteasome inhibitor MG132 or the NEDD8-activating enzyme inhibitor MLN4924. **(A-E)** Vinculin, β-actin, or GAPDH were used as loading controls. Blots are representative of three independent experiments.

We next investigated whether K48-linked polyubiquitination, a canonical signal for proteasomal degradation, enhances proteasome-dependent protein degradation. To this end, Jurkat cells were treated with FK506 in the presence or absence of the proteasome inhibitor MG132. Co-treatment with MG132 led to a marked accumulation of K48-linked polyubiquitin chains compared to FK506 treatment alone, consistent with proteasome-dependent turnover. The observed ubiquitination was sensitive to MLN4924, a potent and selective inhibitor of the NEDD8-activating enzyme that broadly suppresses Cullin ligase activity, indicating that Cullin-RING ligases contribute to the ubiquitination response. Notably, one potential mediator is the CUL4-DDB1 ligase adaptor AMBRA1, which emerged as a strong hit in the CRISPR screen (Soucy *et al*., 2009) (Figure 2E).

Given that HECTD1 and HECT-like TRIP12, E3 ligases identified as top hits in the screen, are reported to catalyse K29-linked ubiquitin chains and ZRANB1/TRABID is a K29/33-specific DUB, we asked whether FK506 also induces K29 ubiquitination. Consistent with this hypothesis, FK506 treatment increased cellular K29-linked polyubiquitin with kinetics similar to the K48 response, while total ubiquitin levels remained unchanged (Figure S2B-C). Consistent with the data presented in Figure 2C, a similar behaviour of K29-linked ubiquitination was observed upon treatment with DMH1, as well as with the combined DMH1/FK506 treatment (Figure S2D).

In summary, these data indicate that FK506 induces a BMP-dependent remodelling of the ubiquitination landscape and that this response is contingent on the activation status of T cells, highlighting a context-dependent coupling between BMP signalling and ubiquitin-mediated regulation in T cell activation.

### BMP-ubiquitin signalling modulates activation responses in primary human T cells

To determine whether the BMP-ubiquitin axis uncovered in Jurkat cells functionally impacts primary T cell responses, we asked if calcineurin-dependent T cell activation is modulated by engagement of this pathway. We thus turned to primary T cells isolated from healthy donors, which allow experimentally controlled activation, particularly important in light of our previous findings showing that ubiquitin accumulation occurs predominantly in activated cells. Using a panel of pharmacological tools to selectively modulate calcineurin activity, BMP receptor signalling, and ubiquitin machinery components, we investigated the functional interplay between these molecular networks during T cell activation. We quantified activation by measuring surface expression of CD154 (CD40L) as a functional readout (Ford *et al*., 1999).

Consistent with its established immunosuppressive activity, treatment with FK506 markedly reduced PMA/I-induced CD154 upregulation in primary human T cells (Figure 3A and S3A). Similarly, inhibition of BMP type I receptor with DMH1 diminished CD154 expression, indicating that endogenous BMP signalling contributes positively to activation upon calcineurin engagement (Figure 3B). Importantly, the degree of suppression observed with DMH1 was comparable to that seen with FK506. Co-treatment with FK506 and DMH1 further reduced CD154 expression compared to both compounds alone, supporting the notion that BMP signalling provides a positive modulation of T cell activation that is independent of calcineurin inhibition (Figure 3C). In line with this, degradation of FKBP12 using 5a1, a selective FKBP12-targeting Proteolysis Targeting Chimera (PROTAC), resulted in a significant increase in CD154 expression upon stimulation (Figure 3D) (Geiger *et al*., 2024). Further, the calcineurin-sparing FKBP12 ligand 18^(S)-Me^, which displaces FKBP12 from BMP receptors without inhibiting calcineurin, was also able to restore T cell activation to normal values (Figure 3E) (Kolos *et al*., 2021). Notably, both 5a1 and 18^(S)-Me^ enhanced T cell activation not only by reversing FK506-induced inhibition, but also by enhancing ongoing calcineurin-dependent responses (Figure 3F-G). This effect was restricted to cells previously stimulated with PMA/I, revealing a relay mechanism in which BMP pathway activation amplifies calcineurin-driven signalling (S3B-C).

**Figure 3.**
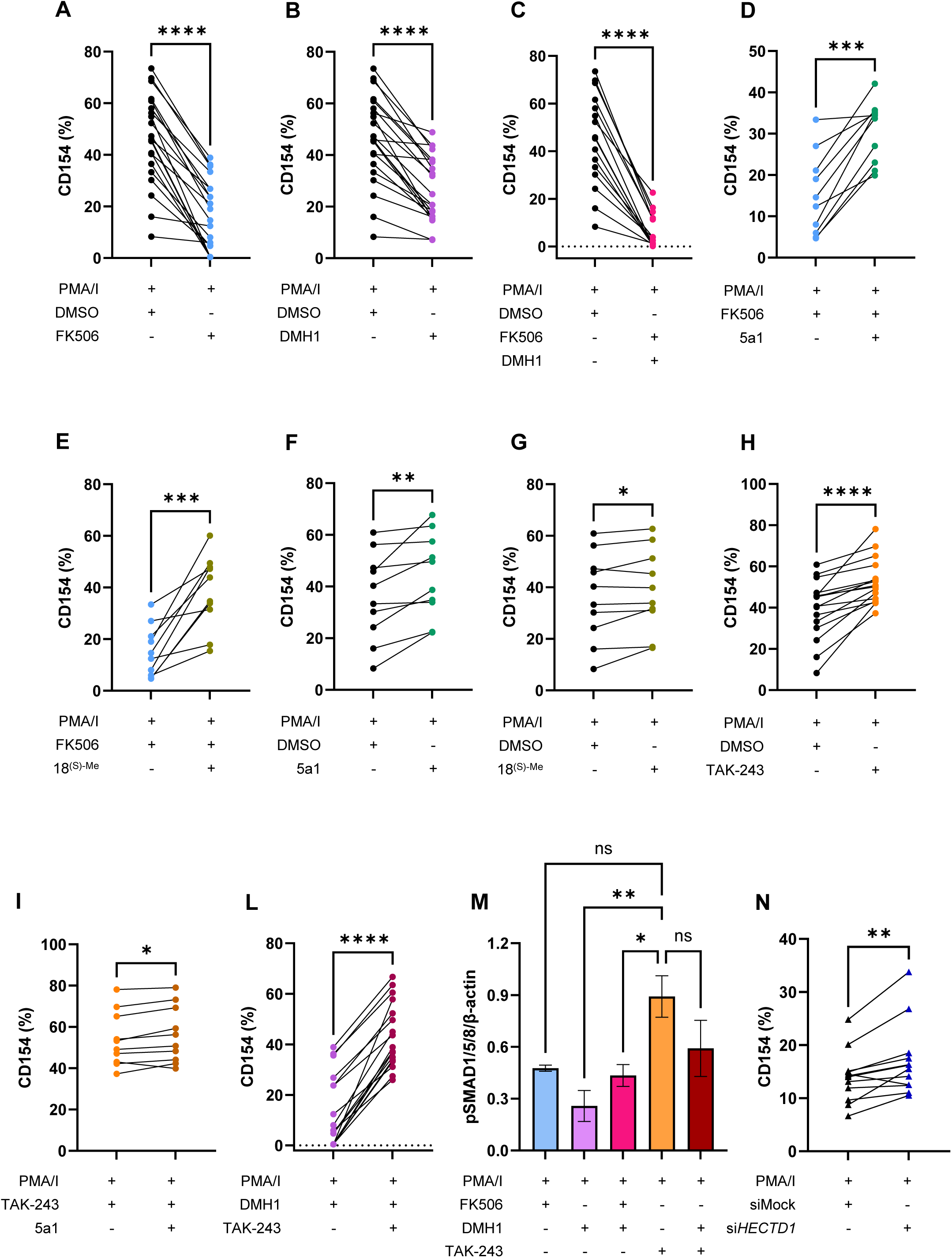
BMP-ubiquitin signalling modulates activation responses in primary human T cells. (A-L) Quantification by flow-cytometry of CD154 surface expression in primary human T cells stimulated with PMA/Ionomycin (PMA/I) in the presence or absence of FK506 (A), BMP type I receptor inhibitor DMH1 (B), FK506 and DMH1 (C), FKBP12-targeting PROTAC 5a1 alone or in combination with FK506 (D), the calcineurin-sparing FKBP12 ligand 18^(S)-Me^ alone or in combination with FK506 (E), 5a1 or 18^(S)-Me^, respectively, during PMA/I stimulation (F-G), the ubiquitin-activating enzyme 1 inhibitor TAK-243 during PMA/I stimulation (H), 5a1 alone or in combination with TAK-243 (I), and TAK-243 alone or in combination with DMH1 (L), as indicated. **(M)** Western blot quantification of pSMAD1/5/8 protein expression relative to β-actin in primary human T cells treated with PMA/I, FK506, DMH1, and TAK-243 alone or in combination, as indicated. Data are presented as mean ± SD from three independent healthy donors. **(N)** Quantification by flow-cytometry of CD154 expression in primary human T cells transfected with control siRNA (siMock) or si*HECTD1* and stimulated with PMA/I. HECTD1 depletion was confirmed by Western blotting (Figure S3F). **(A-L and N)** Each dot (A-L) or triangle (N) represents a single healthy T cell donor. **(A-N)** Statistical significance was assessed by paired t-test. ns, not significant; *, p < 0.05; **, p < 0.005; ***, p < 0.0005; ****, p < 0.0001.

We next asked whether ubiquitin-dependent processes functionally intersect with BMP signalling during T cell activation. Our CRISPR screen identified components of the ubiquitin machinery, including the E2 conjugating enzyme UBE2K and the HECT-type E3 ligase HECTD1, as negative regulators of FK506 response, suggesting that ubiquitin conjugation restrains BMP-driven effects. Consistently, pharmacological inhibition of the ubiquitin-activating enzyme 1 (UAE1), a central component of ubiquitin conjugation that functionally cooperates with HECT-type E3 ligases such as HECTD1, using TAK-243, enhanced CD154 upregulation upon PMA/I stimulation (Figure 3H) (Bedford *et al*., 2011). Again, this effect was restricted to previously stimulated T cells, with no impact on quiescent T cells, indicating that inhibition of ligases downstream of UAE1 amplifies calcineurin-dependent activation (Figure S3D). Co-treatment with TAK-243 and 5a1 further increased CD154 expression in a synergistic manner, reinforcing the functional interplay between BMP signalling and ubiquitin-dependent regulation in shaping T cell activation (Figure 3I). Importantly, TAK-243 treatment was able to rescue T cell activation after BMP pathway blockade by DMH1 (Figure 3L). Consistent with these findings, TAK-243 induced a marked increase in pSMAD1/5/8 levels, partially restoring the reduction induced by DMH1 to levels comparable to PMA/I and FK506 combined treatment (Figure 3M). Finally, depletion of HECTD1, an E3 ligase downstream of UAE1, significantly increased CD154 expression following PMA/I stimulation, phenocopying TAK-243 treatment and supporting the hypothesis that HECTD1 acts as a negative regulator of T cell activation within the BMP-ubiquitin axis (Figure 3N and S3E-F).

Collectively, these data indicate that BMP receptor signalling and ubiquitin-dependent processes constitute an integrated signalling network that modulates T cell activation. While FK506 suppresses stimulation primarily through calcineurin inhibition, its concomitant induction of BMP signalling engages a ubiquitin-dependent program that supports and counterbalances T cell activation.

### FK506 induces a BMP-associated ubiquitin remodelling and proteasome-dependent turnover of signalling regulators

To identify proteins ubiquitinated in response to BMP pathway activation by FK506, we turned to a proteomics-based approach. We used ubiquitin-affinity nanobody-based beads to capture ubiquitylated proteins in Jurkat cells and isolated ubiquitinated proteins from cells treated with FK506 alone or in combination with MG132, with DMSO and MG132 treatments as controls, allowing the identification of proteins targeted for degradation in response to FK506 (Figure 4A and S4A). Overall, we identified 52,042 unique peptide sequences across the samples, mapping to 5,570 unique proteins, of which 3,254 were consistently detected across replicates and quantifiable (Figure S4B and Source data 1). To identify proteins undergoing proteasomal degradation in response to FK506, we selected candidates that were enriched in the FK506/MG132 condition relative to FK506 treatment alone, while excluding those enriched upon MG132 treatment alone. The remaining proteins are therefore likely to represent substrates specifically targeted for degradation following FK506 treatment. Consistent with this approach, we identified several proteins involved in BMP signalling, including the E3 ligases TRIM33, MEN1, and the SUMO E3 ligase ZNF451 (Figure 4B) (Kaji *et al*., 2001; Feng *et al*., 2014; Guo *et al*., 2017).

**Figure 4.**
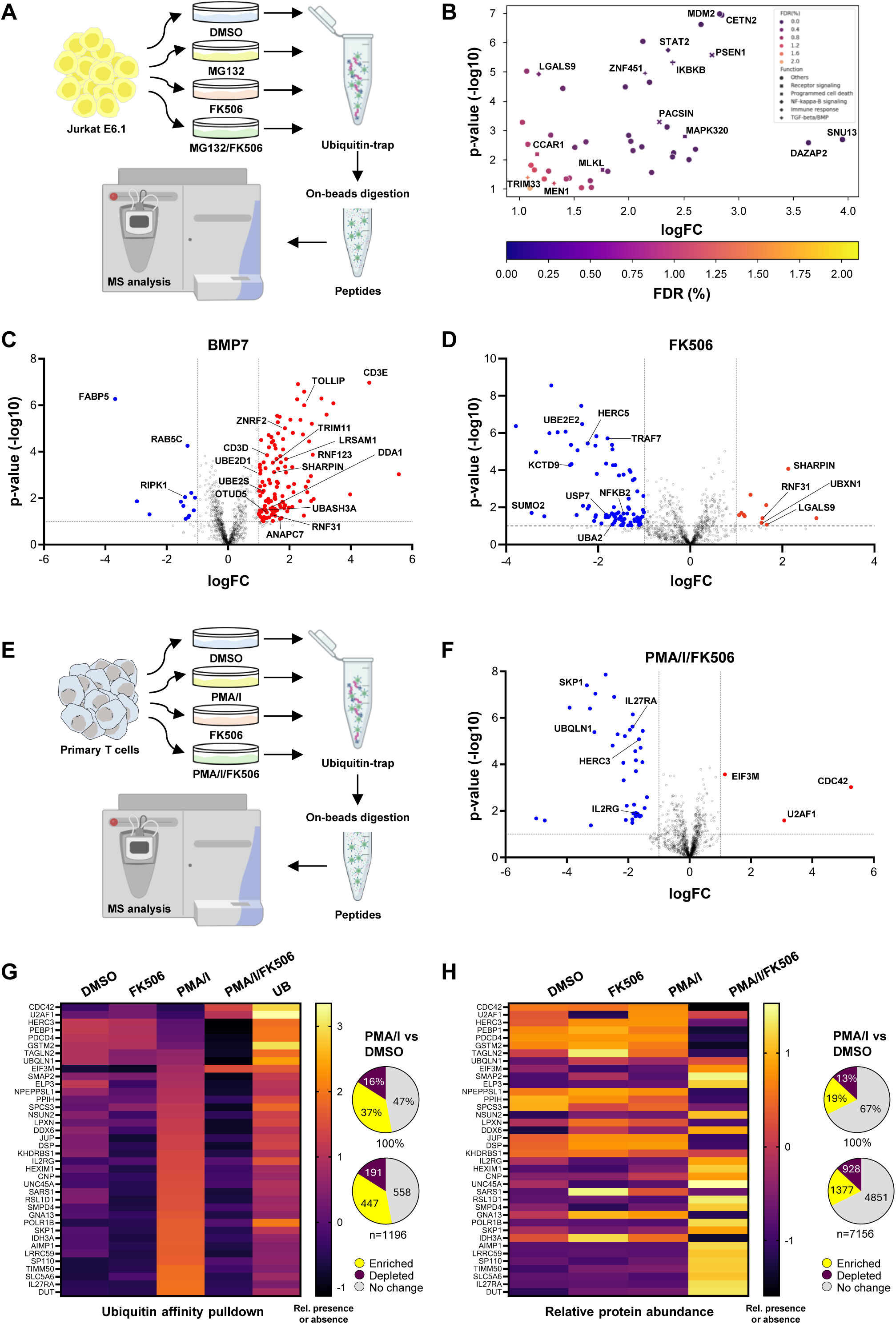
FK506 induces BMP-associated ubiquitin remodelling and proteasome-dependent turnover of signalling regulators. **(A)** Schematic overview of the ubiquitin-enrichment proteomics workflow in Jurkat E6.1 cells. Cells were treated with DMSO, MG132, FK506, or a combination of MG132 and FK506, followed by enrichment of ubiquitinated proteins using ubiquitin-affinity nanobody beads, on-bead digestion, and LC-MS/MS analysis. The mass spectrometry instrument and the reagent tube images were taken from BioRender. **(B)** Differential enrichment plot showing proteins enriched in the MG132/FK506 condition relative to FK506 treatment alone after excluding proteins enriched by MG132 alone. Highlighted proteins represent candidate substrates undergoing proteasome-dependent turnover in response to FK506 treatment. The colour scale indicates the false discovery rate (FDR). **(C-D)** Volcano plots of ubiquitinated proteins identified by ubiquitin-enrichment mass spectrometry in Jurkat E6.1 cells treated with recombinant BMP7 (C) or FK506 (D), relative to DMSO-treated controls. Selected immune-related and ubiquitin-associated proteins are indicated. Differentially enriched or depleted proteins are highlighted in red and blue, respectively. Dashed lines indicate significance and fold-change thresholds. **(E)** Schematic overview of the ubiquitin-enrichment proteomics workflow in primary human T cells. Naïve or PMA/Ionomycin (PMA/I)-activated T cells were treated with FK506 as indicated, followed by ubiquitin-affinity enrichment and LC-MS/MS analysis with the same approach in Figure 4A. **(F)** Volcano plot showing ubiquitinated proteins differentially regulated in PMA/I/FK506-treated primary human T cells compared to activated T cells (PMA/I-treated). Selected immune-related and ubiquitin-associated proteins are indicated. Differentially enriched or depleted proteins are highlighted in red and blue, respectively. Dashed lines indicate significance and fold-change thresholds. **(G)** Heatmap representation of ubiquitinated proteins differentially regulated across DMSO, FK506, PMA/I, and PMA/I/FK506 conditions in primary human T cells. Pie charts summarize the proportion of enriched, depleted, and unchanged ubiquitinated proteins in PMA/I-treated cells relative to DMSO controls. UB, ubiquitin pulldown enrichment score. **(H)** Heatmap representation of relative protein abundance across DMSO, FK506, PMA/I, and PMA/I/FK506 conditions in primary human T cells. Pie charts summarize the proportion of proteins significantly enriched, depleted, or unchanged in PMA/I-treated cells relative to DMSO controls. **(G-H)** Colour scale represents log2 fold change in ubiquitination abundance (G) and relative protein abundance (H), with yellow indicating increased and deep purple reduced values relative to control.

Next, we sought to assess changes in the ubiquitinome following BMP pathway activation and to compare these with those observed upon FK506 treatment. To this end, we performed ubiquitin-enrichment mass spectrometry in Jurkat cells treated with recombinant BMP7, FK506, or DMSO. In total, we quantified 1,528 proteins, of which 1,101 were putative ubiquitination targets. Relative to DMSO-treated controls, 169 proteins were differentially enriched upon BMP7 treatment, whereas 146 proteins were differentially enriched following FK506 treatment (Figure S4C and Source data 2). BMP7 stimulation induced selective changes in protein ubiquitination: key T cell signalling components, including the T cell receptor (TCR) subunits CD3D and CD3E, as well as UBASH3A, a known positive regulator of TCR accumulation, showed increased levels of ubiquitination. In addition, multiple E3 ubiquitin ligases (ZNRF2, TRIM11, LRSAM1, RNF13, ANAPC7), two E2 ubiquitin-conjugating enzymes (UBE2D1 and UBE2S) and the deubiquitinase OTUD5, previously identified in our CRISPR screen, displayed increased ubiquitination, as well as TOLLIP and DDA1, two ubiquitin adaptor proteins. Notably, components of the LUBAC complex (SHARPIN and RNF31), which mediates linear ubiquitin chain assembly and regulates NF-κB signalling through inhibition of RIPK1, were also enriched, whereas RIPK1 showed decreased ubiquitination (Figure 4C) (Ishikawa *et al*., 2006; Jin *et al*., 2008; Hoxhaj *et al*., 2012; Huett *et al*., 2012; Lu *et al*., 2014; Shabek *et al*., 2018; Ge *et al*., 2019; Zhang *et al*., 2023). These findings indicate that BMP signalling drives a selective reprogramming of the ubiquitinome rather than a global ubiquitin shift. FK506 also promoted selective ubiquitination of key signalling proteins, with overlapping candidates with BMP7 treatment such as SHARPIN and RNF31 (components of the LUBAC complex), supporting the idea that FK506 engages BMP-related ubiquitin remodelling (Figure 4D). A direct comparison of BMP7- and FK506-induced ubiquitination revealed both shared and treatment-specific targets (Figure S4D). While a subset of 24 proteins was commonly regulated by both stimuli, FK506 uniquely affected additional factors, including UBXN1, the E3 ubiquitin ligases HERC5 and TRAF7, and other E2 ubiquitin conjugating and deubiquitinating enzymes, suggesting partial overlap but also distinct ubiquitin remodelling outcomes (Gao *et al*., 2023; Sinha *et al*., 2026). Together, these findings support the conclusion that BMP signalling and FK506 treatment converge on a common ubiquitin-dependent regulatory module, while also activating distinct branches of ubiquitin signalling that may reflect pathway-specific or context-dependent responses (Figure S4E).

We next asked whether the ubiquitinome changes observed upon FK506 treatment are influenced by the activation state of T cells. To address this, primary human T cells isolated from healthy donors were either maintained in a naïve state or treated with PMA/I. Activated cells were then either maintained as untreated controls or treated with FK506. Ubiquitinated proteins from each condition were enriched using ubiquitin-affinity nanobody beads and analysed by quantitative mass spectrometry (Figure 4E). Across all conditions, we quantified 1,411 proteins, of which 1,208 were classified as putative ubiquitin substrates. Global analysis revealed that T cell activation alone induced extensive remodelling of the ubiquitinome. Out of 1,196 identified proteins, 447 were significantly enriched and 191 were depleted in PMA/I-treated cells compared to naïve controls, while 558 proteins remained unchanged (Figure S4F and Source data 3). Thus, more than 50% of the detected ubiquitinome was differentially regulated upon T cell activation, highlighting ubiquitination as a major regulatory change during this transition. Importantly, FK506 treatment of stimulated T cells partially reversed the activation-induced ubiquitination changes. Specifically, 88 proteins that were differentially ubiquitinated upon activation displayed a shift back toward basal ubiquitination levels following FK506 treatment (Figure 4F). Among these were key immune regulators, including the interleukin receptor subunits IL2RG and IL27RA, which mediate cytokine signalling pathways that promote T cell activation, as well as ubiquitin-associated proteins such as UBQLN1, the SCF complex adaptor SKP1, and the E3 ubiquitin ligase HERC3 (Salcedo *et al*., 2009; Whiteley *et al*., 2017; Hong *et al*., 2024; Sinha *et al*., 2026). These findings suggest that FK506 can selectively counteract activation-induced ubiquitination in T cells. In contrast, FK506 treatment in previously stimulated T cells can also induce unique ubiquitination changes not observed under either condition alone, further supporting a context-dependent remodelling of the ubiquitinome (Figure 4G). To distinguish between proteolytic and non-proteolytic ubiquitination events, we performed an analysis of the total proteome. Across conditions, 7,163 proteins were reliably quantified, revealing that PMA/I stimulation altered the abundance of approximately 30% of the proteome. Notably, several proteins associated with ubiquitin-dependent (SKP1 and UBQLN1) and immune processes (IL2RG and IL27RA) exhibited reduced ubiquitination upon FK506 co-treatment, accompanied by a corresponding increase in protein abundance. This inverse relationship is consistent with a model in which ubiquitination targets these proteins for proteasomal degradation. However, for many ubiquitinated proteins identified in the enrichment analysis, changes in ubiquitination were not accompanied by changes in protein abundance, indicating that a substantial fraction of the observed ubiquitination events likely reflects non-proteolytic functions. Conversely, other proteins, such as the E3 ubiquitin ligase HERC3, displayed concomitant reductions in both ubiquitination and protein abundance, suggesting that ubiquitination may also exert non-degradative effects that contribute to protein stabilization (Figure 4H and Source data 4).

In summary, our proteomic analysis demonstrates that the transition from naïve to activated T cells is accompanied by widespread remodelling of the ubiquitinome. FK506 selectively reshapes this landscape in activated cells, both reversing activation-induced ubiquitination of specific targets and introducing additional context-dependent modifications. These effects correlate with BMP pathway engagement and support a model in which calcineurin-BMP crosstalk modulates ubiquitin-dependent regulatory programs during T cell activation.

## DISCUSSION

In the present study, we identified a BMP-associated ubiquitin signalling network that modulates calcineurin-dependent activation responses in human T cells. Our comparative CRISPR screens revealed fundamentally distinct genetic dependency profiles of FK506 and CsA, two calcineurin-targeting compounds. In particular, the FK506 screen identified several BMP pathway components, whose depletion conferred a selective fitness advantage under FK506 treatment, supporting the idea that FK506 engages BMP signalling in T cells similarly to what has been previously reported (Spiekerkoetter *et al*., 2013; Peiffer *et al*., 2019; Larraufie *et al*., 2021). The observation that inhibition of BMP receptors reduced T cell activation in primary cells is also consistent with previous reports suggesting that canonical BMP signalling positively regulates human CD4⁺ T cell activation, proliferation and IL-2 production (Martínez *et al*., 2015). Our data extend these observations by suggesting that the BMP pathway is part of a signalling relay downstream of calcineurin. The chemogenetic profile of FK506 further showed a strong enrichment of ubiquitin-related genes and suggested that ubiquitin-dependent mechanisms are tightly linked to BMP pathway activity in T cells. Consistent with this, FK506 treatment induced rapid accumulation of K48- and K29-linked ubiquitin chains in a BMP receptor-dependent manner. Importantly, this effect was largely restricted to activated primary T cells, indicating that the BMP-ubiquitin axis operates in a context-dependent manner and requires previous T cell activation. Jurkat cells, which display constitutive signalling activity, bypassed this requirement and showed ubiquitin remodelling after FK506 treatment even in the absence of stimulation. Our ubiquitinome analyses further revealed that both FK506 and BMP7 induce selective, rather than global ubiquitin modifications. The observation that proteasome inhibition led to accumulation of FK506-induced ubiquitinated proteins suggests that at least a subset of these events is linked to proteasome-dependent turnover. Interestingly, many ubiquitination changes occurred without detectable alterations in protein abundance, arguing that non-proteolytic ubiquitination also contributes substantially to this response. Notably, several ubiquitin regulators identified in the CRISPR screen were also independently recovered in the ubiquitin proteomics, further supporting a strong concordance between genetic dependencies and dynamic ubiquitin regulation after BMP pathway activation. Functionally, our data provide evidence that BMP signalling positively modulates calcineurin-dependent T cell activation. While FK506 inhibits activation primarily through calcineurin blockade, selective activation of BMP signalling by FKBP12 degradation or calcineurin-sparing FKBP12 ligands enhanced T cell activation in stimulated primary cells. Likewise, inhibition of ubiquitin conjugation using TAK-243 potentiated activation responses and partially rescued the effects of BMP pathway inhibition. Several aspects of this study remain to be clarified. First, although our data establish a strong correlation between BMP signalling and ubiquitin remodelling, the role of individual substrates and ubiquitin ligases responsible for these effects remains incompletely defined. HECTD1 emerged as an attractive candidate, whose depletion was able to restore T cell activation, thus corroborating its scoring as resistance gene in the screen and suggesting a potential role of HECT E3 ubiquitin ligases as pivotal regulators within this axis. Second, our proteomic analyses were performed at the level of ubiquitinated proteins rather than individual ubiquitination sites, limiting mechanistic interpretation of specific modifications. Future studies combining site-resolved ubiquitinomics with genetic perturbation approaches will be important to define the functional relevance of these ubiquitination events. While our study focused primarily on activation responses in Jurkat and primary human T cells, it will be of particular interest to understand how this relay impacts long-term T cell differentiation and exhaustion, especially in light of recent reports linking BMP signalling to T cell dysfunction and responsiveness to immunotherapy (Shahinfar *et al*., 2024).

Collectively, our work identifies a previously underrecognized connection between calcineurin signalling, BMP pathway, and ubiquitin-dependent regulation in T cells. These findings support a model in which BMP signalling and the ubiquitin system are integrated into a context-dependent regulatory network that fine-tunes calcineurin activity and T cell activation.

## MATERIALS AND METHODS

### Compounds

The following compounds were used in the study at the indicated concentration: Ionomycin, 5µg/mL, Santa Cruz Biotechnology (Cat. sc-3592); PMA (Phorbol 12-myristate 13-acetate), 0.5 µg/mL, Sigma-Aldrich (Cat. P1585); FK-506, 20 nM (LD20) for CRISPR-screen and 0.625 nM for the other experiments, Sigma-Aldrich (Cat. PHR1809); Cyclosporin A, 3 µM, Sigma-Aldrich (Cat. 32425); DMH1, 5 µM, Selleck Chem, (Cat. S7146); Recombinant human BMP-7 protein, 100 nM, Novus Biological (Cat. 354-BP-010); MG-132, 2 µM, Med Chem Express (Cat. HY-13259); TAK-243, 1 µM, Med Chem Express (Cat. HY-100487); MLN4924, 2.5 µM, Med Chem Express (Cat. HY-70062); FKBP12-PROTAC 5a1, 1µM, and 18^(S)-Me^, 10 µM, were kindly provided by the Hausch Lab, Technical University Darmstadt, Germany.

### Antibodies

Following primary and secondary antibodies were used at the indicated dilutions: Anti-mouse IgG, 1:10,000, HRP-linked Antibody, Cell Signaling (Cat. 7076); Anti-rabbit IgG, 1:10,000, HRP-linked Antibody, Cell Signaling Technology (Cat. 7074); Anti-β-Actin, 1:2,500-1:5,000, Santa Cruz Biotechnology (Cat. sc-47778); Anti-GAPDH, 1:2,000, LICORbio (Cat. 926-42216); Anti-Vinculin, 1:2,000, Sigma-Aldrich (Cat. V9131); Anti-phospho SMAD1/5/8, 1:1,000, Merck (Cat. AB3848-I); Anti-phospho SMAD2, 1:1,000, Cell Signaling Technology (Cat. 3108); Anti-Ubiquitin (P4D1), 1:3,000, Cell Signaling Technology (Cat. 3936). Anti-Ubiquitin K48, 1:1,000, Sigma-Aldrich (Cat. ZRB2150); Anti-Ubiquitin K29, developed in house (*Yu et al*., 2021); Anti-HECTD1, 1:500, Abcam (Cat. Ab101992); Anti-CD3, 1:100, APC-Cyanine7, Biolegend (Cat. 300426); Anti-CD4, 1:50, PE, Biolegend (Cat. 300508); Anti-CD8, 1:50, FITC, Biolegend (Cat. 344704); Anti-CD154, 1:10, APC, BD Biosciences (Cat. 555702).

### Cell culture methods

Primary human T cells (see below) and Jurkat E6.1 cells were cultured at 37°C in 5% CO_2_ humidified environment and maintained in RPMI Medium + GlutaMax supplemented with 10% Fetal Bovine Serum (FBS), Thermo Fisher Scientific (Cat. A5256701) and 1% Penicillin-Streptomycin 10,000 U/ml, Thermo Fisher Scientific (Cat. 15140122). Jurkat E6.1 cells were purchased from ATCC (TIB-152) and were not tested for mycoplasma. Medium used for RNAi experiments was not supplemented with Penicillin-Streptomycin.

### Isolation of primary T cells

Primary T cells were isolated from buffy coats of voluntary healthy donors of the DRK-Blutspendedienst Baden-Württemberg ǀ Hessen in Mannheim, Germany (informed consent was obtained; no specific ethical approval was required). Blood samples were mixed with an equal amount of PBS supplemented with 2 mM EDTA. The blood-PBS mixture was then overlaid over Ficoll (Pancoll human, Cat. P04-60100) and separation of the different blood components was performed by density gradient centrifugation at 440×g for 30 min. After centrifugation, the mononuclear cell fraction was collected and washed twice with 2 mM ETDA in PBS. Cells were then lysed with RBC lysing buffer (PluriSelect, Cat. 60-00050-11) and cleared by centrifugation at 300×g for 10 min. Pellets were washed once with 2 mM EDTA in PBS. Cell pellets were then resuspended in MACS buffer (2 mM EDTA, 0.5% BSA in PBS) and T cells were isolated by negative selection with the Pan T Cell Isolation Kit, human (Miltenyi Biotec, Cat. 130-096-535). Purity of the isolated T cells was assessed by flow-cytometry. Cells were used fresh and were not subjected to freezing at any stage prior to the experiments.

### CRISPR-Cas9 knockout screens

Lentiviral particles were produced in HEK293T cells purchased from ATCC by transfection of the “all-in-one” LCV2::TKOv3 pooled library, Addgene (Cat. 90294) together with the lentiviral packaging plasmid psPAX2, Addgene (Cat. 12260), and the envelope plasmid pMD2.G, Addgene (Cat. 12259), using Lipofectamine 3000, Invitrogen (Cat. L3000) in OptiMEM, Gibco. Six hours post-transfection, the medium was exchanged with RPMI Medium + GlutaMax supplemented with 10% FBS and 1% Penicillin-Streptomycin 10,000 U/ml. 48 h post-transfection, viral particles were harvested and filtered through a 0.45 µm syringe filter and stored at −80°C until use. The LCV2::TKOv3 sgRNA library was a gift from Jason Mofat. pMD2.G and psPAX2 plasmids were gifts from Didier Trono. Screens were performed as duplicates at a coverage of >300-fold sgRNA representation. Jurkat cells were transduced with the library at a low multiplicity of infections (MOI, 0.2-0.3) by incubating cells with 8 µg/mL polybrene together with lentiviral supernatant for 24 h. Transduced cells were selected by treatment with 3.5 µg/mL puromycin for 48 h. After selection, cells were expanded for 6 days and further split into mock-treated, FK506, or Cyclosporin A treated conditions. Cells were expanded for an additional 12 days with passage every 3 days in fresh medium alone or with 20 nM FK506 or 3 µM Cyclosporin A. Pellets were harvested at the start of the screen (T0), and at the final timepoint (T18). Genomic DNA extraction, next-generation sequencing and analysis were performed as previously described (Vit *et al*., 2022).

### Transient siRNA transfection

Transfection of siRNA was performed using the Amaxa human T cell nucleofector kit, Lonza (Cat. VPA-1002), according to the manufacturer’s instructions. RNAi oligonucleotides were purchased by Dharmacon and used as a pool of four targeting sequences: ON-TARGETplus non-targeting pool, Dharmacon (Cat. D-001810-10-05): 1: UGGUUUACAUGUCGAGUAA; 2: UGGUUUACAUGUUGUGUGA; 3: UGGUUUACAUGUUUUCUGA; 4: UGGUUUACAUGUUUUCCUA. ON-TARGETplus human HECTD1 (25831), Dharmacon (Cat. LQ-007188-00-0002): 1: GUUAAUAGCUGUACUAGAA; 2: GCUCAUAGCUGCAUAUAAG; 3: CAUAGAGGAUUUAGGUUUA; 4: GAAAGGGACAUGCAACUAA. Efficient depletion of HECTD1 was assessed by Western blotting.

### Western blotting

Harvested cells were lysed on ice with lysis buffer (50 mM Tris/HCL, pH 7.4, 150 mM NaCl, 1 mM EDTA, 0.25% sodium deoxycholate, 1% NP-40) supplemented with protease inhibitor, cOmplete, Roche, and phosphatase inhibitor, PhosSTOP, Roche. For ubiquitin blots, lysis buffer was additionally supplemented with 1.25 mM N-ethylmaleimide, Sigma (Cat. 04260), and 50 µM PR-619, Sigma (Cat. 662141). After centrifugation for 30 min at 4°C at maximum speed, the supernatant was collected and supplemented with a 1:4 dilution of NuPage LDS Sample Buffer (Invitrogen by Thermo Fisher Scientific) containing 15% 2-Mercaptoethanol. The samples were then heated at 95°C for 7 min. Lysates were resolved in NuPage 10% Bis-Tris Gel, Thermo Fisher Scientific, or in NuPage 4-12% Bis-Tris Gel, Thermo Fisher Scientific. Gel electrophoresis was performed with MOPS SDS Running Buffer, Invitrogen by Thermo Fisher Scientific, and gels were transferred onto mini PVDF Membrane, Biorad, or onto IMMUN-Blot low-fluorescence PVDF membranes, Biorad. Western blot analysis was carried out using standard protocols with primary antibodies and horseradish peroxidase (HRP)-conjugated secondary antibodies (see Antibodies section) and bands were visualized by using SuperSignal West Chemiluminescent Substrate, Thermo Fisher Scientific, and captured using an Odyssey M imager (Li-Cor) or ChemiDoc imaging system, Biorad. Alternatively, IRDye far red secondary antibodies (Li-Cor) were used.

### Flow cytometry

Primary T cells were treated as follows: 0.625 nM FK506 for 6 h; 5 µM DMH1 for 14 h; 1µM TAK-243 for 2 h; 1 µM FKBP12-Protac 5a1 for 8 h; 10 µM 18^(S)-Me^ for 10 min; Ionomycin and PMA were added to the cells simultaneously for 2 h at 5 µg/mL and 0.5 µg/mL, respectively. After treatment, cells were harvested for staining with antibodies (see Antibodies section). T cells were washed with PBS and resuspended in 100 µL of FACS Buffer and 10 µL of FcR blocking reagent, Miltenyi Biotec (Cat. 130-059-901). Antibodies were incubated for 20 min at 4°C. After washing, T cells were resuspended in FACS buffer supplemented with Sytox Blue, 1:2000, Thermo Fisher Scientific (Cat. S34857).

### Ubiquitin affinity pulldown and Mass Spectrometry analysis

#### MS sample preparation

Jurkat E6.1 cells were seeded to a density of 1×10^6^ cells/mL in 50 mL medium medium containing either DMSO, 100 nM recombinant BMP7, or 100 nM FK506 (each condition in quadruplicate) for 2 h. As a technical control, additional DMSO-treated samples were included in quadruplicate. After treatment, cells were collected into 50 mL tubes and centrifuged at 1,000 g at 4 °C, washed once with ice cold PBS containing 10 µM PR-169 and transferred to 10 mL tubes to be centrifuged at 1,000 g at 4°C. Pellets were resuspended in lysis buffer containing 50 mM Tris (pH 7.4), 500 mM NaCl, 1% NP-40, to which protease inhibitor cocktail tablets, phosphatase inhibitor tablets (1 tablet/10 ml each), NEM (1.25 mg/ml), PR-169 (10 µM), and DTT (1 mM) were added fresh. Resuspended cells were incubated on ice for 30 min and then sonicated and cleared by centrifugation at 20,000×g for 30 minutes. Subsequently, lysates were quantified using a Pierce BCA assay and an equal amount of lysate was incubated with ubiquitin-trap agarose beads, ChromoTek, for one hour at 4°C with rotation. Technical control samples were instead incubated with GFP-trap agarose beads, ChromoTek. Beads were then washed three times with lysis buffer, transferred to fresh 1.5 mL tubes, and then washed a further four times with a reduced wash buffer containing 50 mM Tris (pH 8) and 50 mM NaCl.

During the last wash, beads were moved to fresh 1.5 mL tubes and remaining wash buffer was removed using a 27 G syringe. On beads digestion was done by adding two bead-volumes of buffer containing 50 mM Tris (pH 8.5) and 2.5 ng/µl sequencing-grade modified trypsin and incubating beads on ice for one hour, making sure to keep them mixed, before moving them to a dry water bath set to 30°C overnight with shaking. The next day, the bead suspension was filter-cleared through 0.45 µm cut-off spin filters, Costar, and stored at −20°C until further analysis. MultiDSK affinity reagent pulldown was done as described in Liu *et al*., 2024. All samples were reduced and alkylated by adding TCEP and chloroacetamide to 5 mM, and incubated at 30°C for 30 minutes. For total proteome analysis, samples were supplemented with SDS to a final concentration of 2%, and homogenized by heating to 99 °C while shaking at 1,400 rpm. Homogenized samples were processed according to the Protein Aggregate Capture protocol (Batth *et al*., 2019). To this end, samples were supplemented with 10 volumes of acetonitrile to precipitate all proteins, after which 50 µL (dry volume) of magnetic microspheres were added. Samples were briefly mixed, beads were allowed to settle, and afterwards placed in a magnetic rack to fix the beads to the side of the tubes. Beads were subsequently washed twice with 100% acetonitrile and once with 70% ethanol, dried and resuspended in 100 µL of ice-cold 50 mM Tris-HCl, pH 8.5, supplemented with 2.5 ng/µL sequencing-grade modified trypsin. Beads were mixed for 10 min on ice, and moved to 30°C for overnight shaking at 1,250 rpm. All samples were reduced and alkylated by adding TCEP and chloroacetamide to 5 mM, and incubated at 30°C for 30 min.

#### Peptide purification

For desalting and purification of peptides, C18 StageTips were prepared in-house, by layering four plugs of C18 material (Sigma-Aldrich, Empore SPE Disks, C18, 47 mm) per StageTip. Activation of StageTips was performed with 100 μL 100% methanol, followed by equilibration using 100 μL 80% acetonitrile in 0.1% formic acid, and two washes with 100 μL 0.1% formic acid. Samples were acidified to pH < 3 by addition of trifluoroacetic acid to a concentration of 1%, after which they were loaded on StageTips. Subsequently, StageTips were washed twice using 100 μL 0.1% formic acid, after which peptides were eluted using 80 µL 30% acetonitrile in 0.1% formic acid. All fractions were dried to completion using a SpeedVac at 60°C. Dried peptides were dissolved in 20 or 40 μL 0.1% formic acid, for pulldown and total proteome samples, respectively, and stored at −20°C until analysis using mass spectrometry.

#### LC-MS analysis

Four datasets (DS1, DS2, DS3, DS4) were analysed for this project, corresponding to Source Data 1, 2, 3, 4 in the manuscript, and with sample preparation details and individual LC-MS settings outlined below. All samples were analyzed on a Vanquish™ Neo UHPLC system (Thermo) coupled to an Orbitrap™ Astral™ mass spectrometer (Thermo). Samples were analyzed on 20 cm long analytical columns, with an internal diameter of 75 μm, packed in-house using ReproSil-Pur 120 C18-AQ 1.9 µm particles (Dr. Maisch). The analytical column was heated to 40 °C, and elution of peptides from the column was achieved by application of gradients with buffer A (0.1% FA) and increasing amounts of buffer B (80% acetonitrile in 0.1% FA). The primary analytical gradient ranged from 8% to 33% buffer B for DS1, 8% to 36% buffer B for DS2, 7% to 35% buffer B for DS3, and 8% to 32% buffer B for DS4. Total gradient length including ramp-up and wash-out was 40 min for pulldown analyses (DS1, DS2, DS4) and 20 min for total proteome analysis (DS3). Primary analytical gradient length was 25 min, 28.5 min, 18 min, and 28.5 min for DS1, DS2, DS3, and DS4, respectively. Ionization was achieved using a NanoSpray Flex™ NG ion source (Thermo). Spray voltage set at 2 kV, ion transfer tube temperature to 275°C, and RF lens to 50%. All full precursor (MS1) scans were acquired using the Orbitrap™ mass analyzer, while all tandem fragment scans were acquired in parallel using the Astral™ mass analyzer. Pulldown samples (DS1, DS2, DS4) were acquired in data-dependent mode (DDA), whereas total proteome samples (DS3) were acquired in data-independent (DIA) mode. For DDA analysis, full scan range was set to 300-1,300 m/z, MS1 resolution to 120,000 (DS1) or 180,000 (DS2, DS4), MS1 AGC target to “500” (5,000,000 charges; DS2, DS4) or “250” (2,500,000 charges; DS1), and MS1 maximum injection time to 150 ms. Precursors with charges 2-6 selected for fragmentation using an isolation width of 1.3 m/z and fragmented using higher-energy collision disassociation (HCD) with normalized collision energy of 25. Monoisotopic Precursor Selection (MIPS) was enabled in “Peptide” mode. Repeated sequencing of precursors was minimized by setting expected peak width to 10 s, and dynamic exclusion duration to 10 s (DS1) or 15 s (DS2, DS4), with an exclusion mass tolerance of 10 ppm and exclusion of isotopes. MS2 fragment scan range was set to 100-1,500 m/z, MS2 AGC target to “100” (10,000 charges), MS2 intensity threshold to 25,000 (DS1) or 50,000 (DS2, DS4) charges per second, and MS2 maximum injection time to 20 ms. Cycle time was set to 0.3 s (DS1) or 0.5 s (DS2, DS4). For DIA analysis, settings were as above with the following adjustments. Full scan range was set to 300-1,000 m/z, and MS1 maximum injection time to 50 ms. Precursors were isolated from the 300-1,000 m/z range, using 4 Th windows (i.e. 175 windows in total), a maximum injection time of 5 ms, and an AGC target of “200” (20,000 charges). Fragmentation was performed using HCD at 25 NCE, with a fragment scan range of 100-1,500 m/z. Loop control was set to “All”.

#### MS data analysis

For pulldown samples (DS1, DS2, DS4), all RAW files were analysed using MaxQuant software v2.5.2.0. (Cox and Mann, 2008; Cox et al., 2011). Default MaxQuant settings were used, with exceptions specified below. For generation of the theoretical spectral library, a HUMAN.fasta database was downloaded from UniProt on the 29th of April, 2023. In-silico digestion of proteins to generate theoretical peptides was performed with trypsin, with maximum number of missed cleavages set to 3. Allowed variable modifications were oxidation of methionine (default), protein N-terminal acetylation (default), deamidation of glutamine and asparagine, peptide N-terminal conversion of glutamine to pyroglutamate, and GlyGly (ubiquitin remnant) on lysine, with maximum variable modifications per peptide set to 3. First search mass tolerance was set to 15 ppm, with main search mass tolerance at 4.5 ppm (default). Second peptide search was disabled. Matching between runs (MBR) was enabled with an alignment window of 10 min and a matching time window of 0.5 min. Label-free quantification (LFQ) was enabled, with “Fast LFQ” disabled. Stringent MaxQuant 1% FDR data filtering at the PSM- and protein-levels was applied (default). For total proteome samples (DS3), analysis of the mass spectrometry raw data was performed using DIA-NN (Demichev *et al*., 2020), v1.9.2. For library-free search, the human FASTA database (105,529 entries) was downloaded from UniProt on the 13th of November, 2024, and DIA-NN was used to generate a spectral library in silico. To this end, settings used were Trypsin/P digestion (default), 2 missed cleavages, peptide length between 6 and 60, precursor charge state between 2 and 4, precursor m/z range from 300 to 1,000 m/z, and fragment ion m/z range from 100 to 1,500 m/z. RAW files were first converted to DIA format, after which they were searched with DIA-NN using the spectral library described above, with default settings. “MBR” was enabled, which generates a new spectral library based on a first search, which is then used to re-analyze the data. Global false discovery rate control was left at 1% in the GUI (default). In addition, the “--matrix-spec-q” and “--matrix-qvalue 0.002” commands were provided, to enforce a post-processing 0.2% FDR control at the run-specific level (in addition to global FDR control at 0.2%).

## DATA AVAILABILITY

The mass spectrometry proteomics data have been deposited to the ProteomeXchange Consortium via the PRIDE [PubMed ID: 39494541] partner repository with the dataset identifier PXD080599.

## STATISTICAL ANALYSIS

Statistical analyses were performed using GraphPad Prism (version 10, GraphPad Software). Details of the statistical tests used for each analysis are provided in the corresponding figure legends.

## ACKNOWLEDGEMENTS

Work at the Novo Nordisk Foundation Center for Protein Research is supported by NNF14CC0001. Work in the Nilsson lab is supported by grants from the Novo Nordisk Foundation (NNF0082227 and NNF0065098) and the Danish Cancer Society (R269-A15586-B71). Work in the Vit lab is supported by grants BSD117779, BSD117781, and BSD117775. We thank Anke Diehlmann and Christina Schmuttermaier for assistance with technical, logistical and organizational aspects of the work. We thank Prof. Felix Hausch for kindly gifting us with the FKBP12-PROTAC 5a1 and 18^(S-Me)^. We also thank the protein production and characterization facility at Novo Nordisk Foundation Center for Protein Research and Center for Stem Cell Medicine for technical support and use of instruments. NGS data processing and analysis were performed using the DeiC National Life Science Supercomputer at Technical University of Denmark (www.computerome.dk).

## AUTHOR CONTRIBUTIONS

**Marcel Diallo**: Conceptualization; Formal Analysis; Validation; Investigation; Visualization; Methodology; Writing – original draft; Writing – review and editing. **Leon Müller**: Conceptualization; Formal Analysis; Validation; Investigation; Visualization; Methodology; Writing – original draft; Writing – review and editing. **Stefanie Uhlig**: Formal Analysis; Validation; Investigation; Methodology. **Ivo Hendriks**: Formal Analysis; Validation; Investigation; Methodology; **Julia Kzhyshkowska**: Critical feedback. **Harald Klüter**: Critical feedback. **Michael Nielsen**: Critical feedback. **Jesper Olsen**: Critical feedback. **Patrick Wuchter**: Critical feedback; Data discussion. **Karen Bieback**: Formal Analysis; Methodology; Critical feedback; Data discussion; Writing – review and editing. **Jakob Nilsson**: Conceptualization; Supervision; Funding acquisition; Methodology; Writing – original draft; Writing – review and editing. **Gianmatteo Vit**: Conceptualization; Supervision; Funding acquisition; Investigation (CRISPR screening); Formal analysis; Visualization; Methodology; Writing – original draft; Writing – review and editing.

---

**Figure S1 (related to Figure 1).**
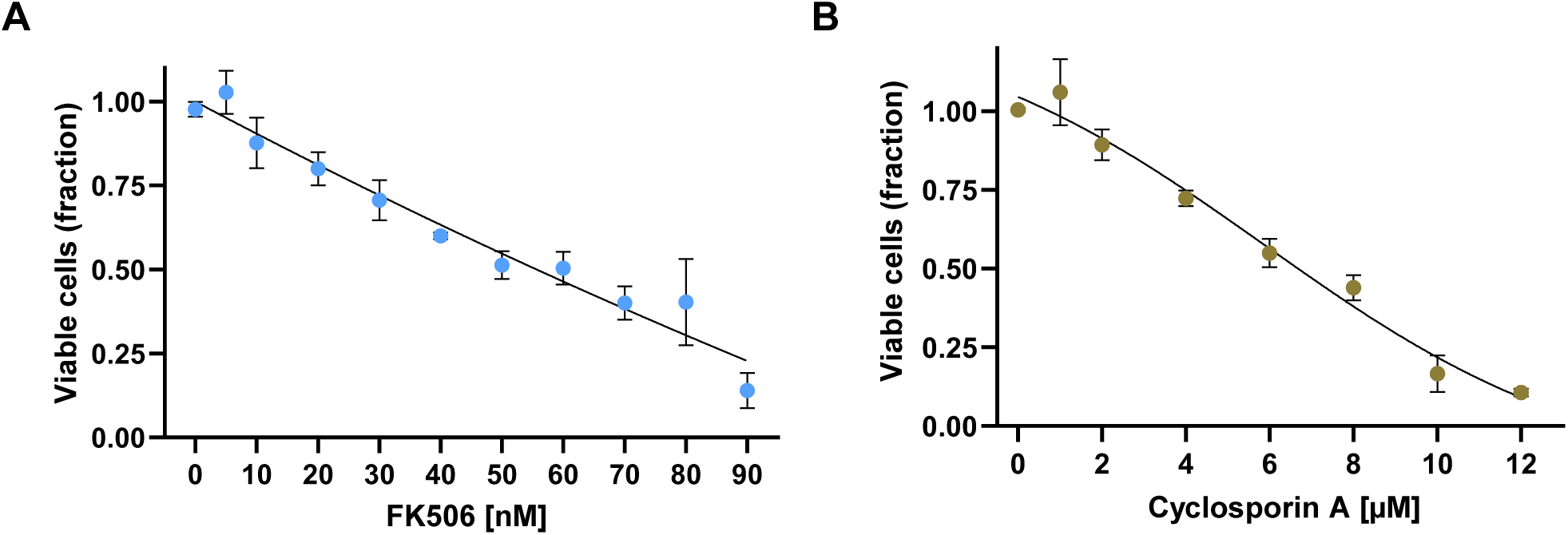
**(A-B)** Determination of LD20 values for FK506 and Cyclosporin A in Jurkat E6.1 cells. Dose-response curves were used to determine the LD20 of FK506 (20 nM) (A) and Cyclosporin A (3 µM) (B). Jurkat E6.1 cells were treated for 12 days with increasing concentrations of the indicated compounds, and cell viability was measured at the end of the treatment period. Viability values are expressed as decimal fractions relative to untreated controls. Values are representative of three independent experiments.

**Figure S2 (related to Figure 2).**
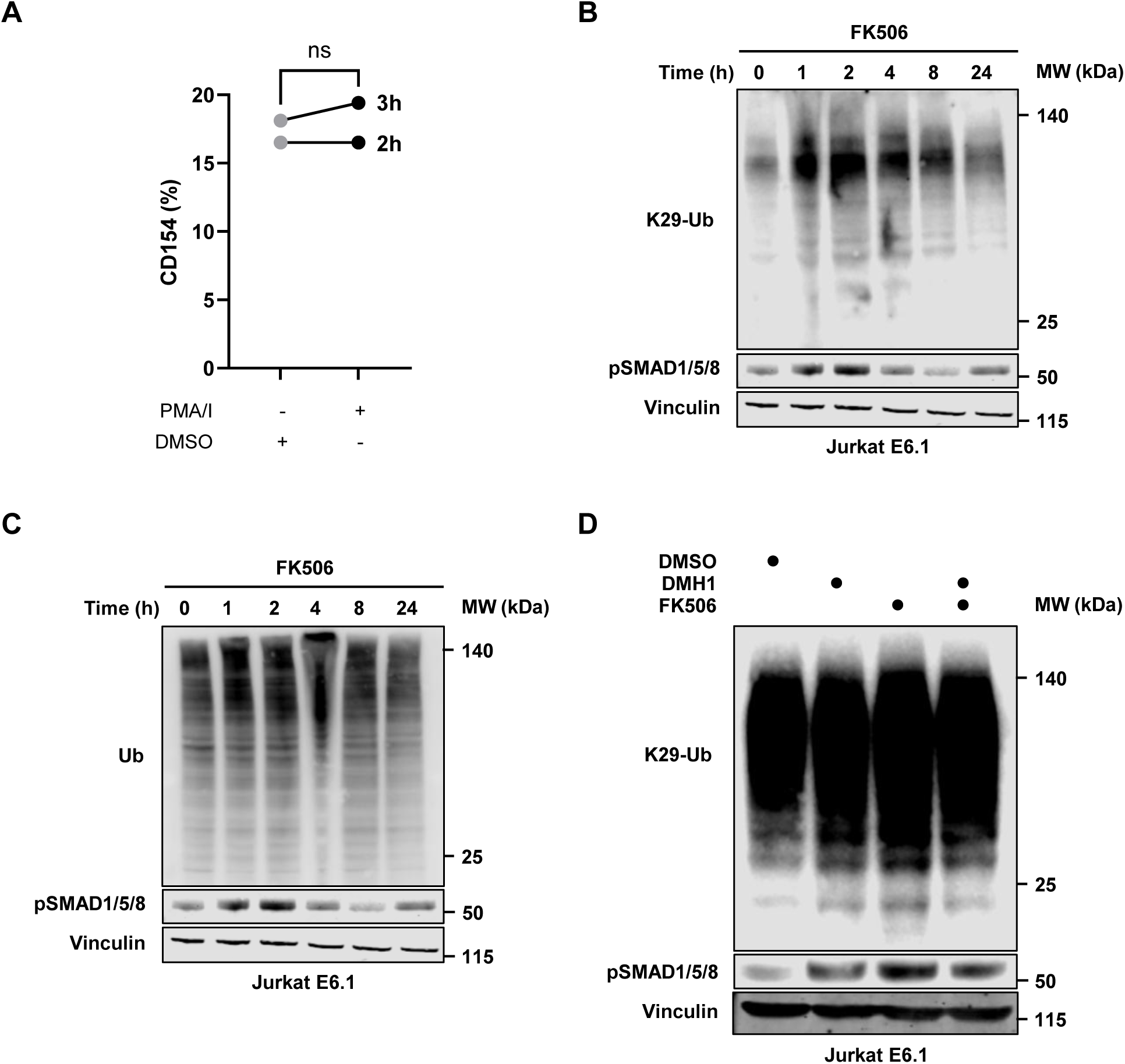
**(A)** Quantification by flow-cytometry of CD154 surface expression in Jurkat E6.1 cells unstimulated or stimulated with PMA/Ionomycin (PMA/I) after two and three hours post activation, as indicated. **(B)** Time-course analysis of K29-linked polyubiquitin chains (K29-Ub) and pSMAD1/5/8 in Jurkat E6.1 cells treated with FK506 at the indicated time points. **(C)** Time-course analysis of total ubiquitin (Ub) levels and pSMAD1/5/8 in Jurkat E6.1 cells treated with FK506 at the indicated time points. **(D)** K29-Ub and pSMAD1/5/8 levels in Jurkat E6.1 cells treated with FK506 or DMH1, alone or in combination, as indicated. (B-D) Vinculin was used as loading control. Blots are representative of three independent experiments.

**Figure S3 (related to Figure 3).**
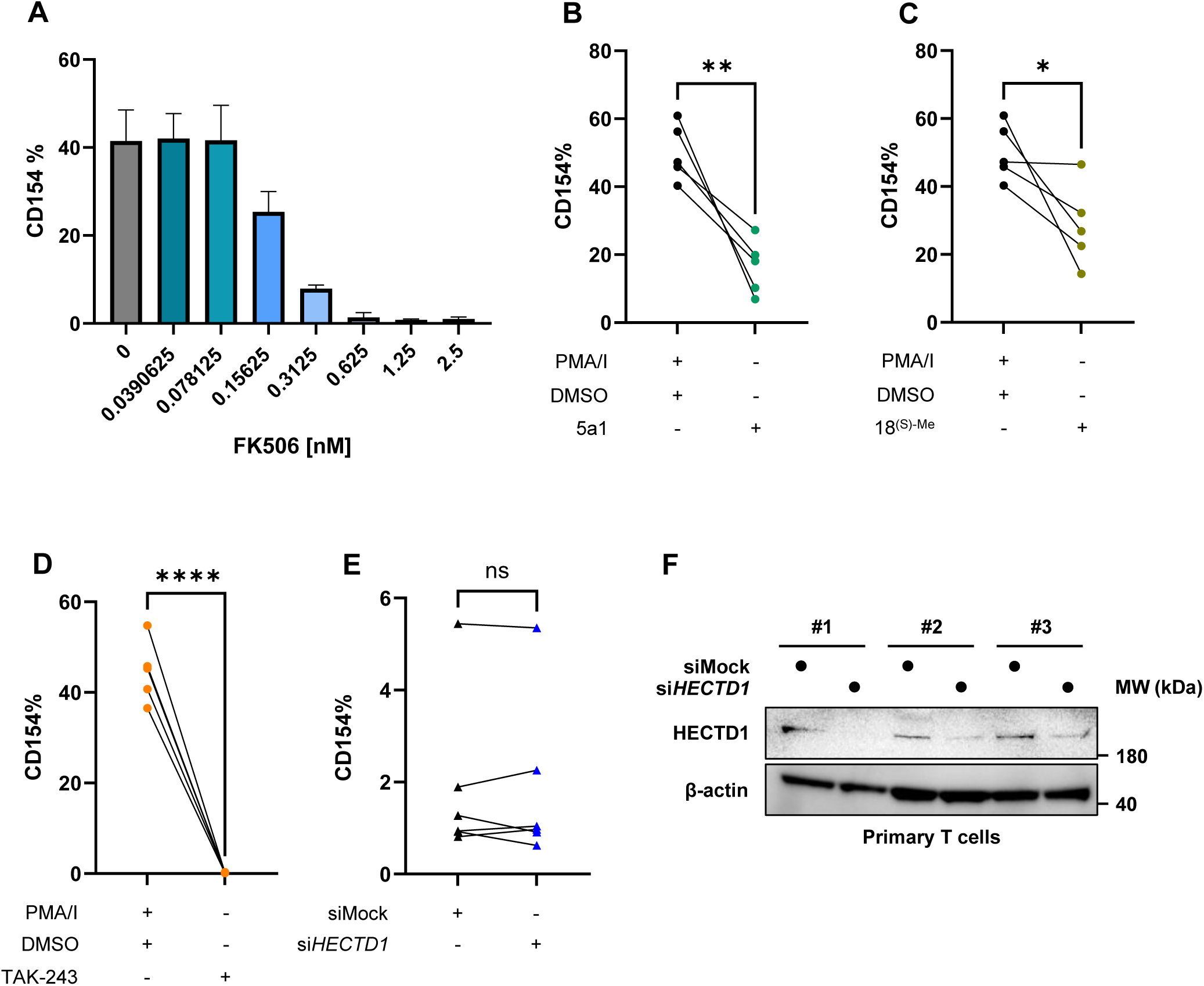
**(A)** Titration of FK506 in activated primary human T cells to determine the optimal concentration that partially suppress T cell activation while preserving a dynamic range of CD154 expression suitable for assessing modulatory effects of additional treatments in downstream functional assays. Primary human T cells were stimulated with PMA/Ionomycin (PMA/I) and treated with increasing concentrations of FK506, as indicated. T cell activation was quantified by flow-cytometric analysis of CD154 surface expression. Data are presented as mean ± SD from three independent healthy donors. **(B-D)** Quantification by flow-cytometry of CD154 surface expression in primary human T cells after treatment with FKBP12-targeting PROTAC 5a1 (B), the calcineurin-sparing FKBP12 ligand 18^(S)-Me^ (C), or the ubiquitin-activating enzyme 1 inhibitor TAK-243 (D), with or without PMA/I stimulation (B-D). **(E)** Quantification by flow-cytometry of CD154 expression in primary human T cells transfected with control siRNA (siMock) or si*HECTD1* without previous stimulation with PMA/I. HECTD1 depletion was confirmed by Western blotting (Figure S3F). **(B-E)** Each dot (B-D) or triangle (E) represents a single T cell healthy donor. Data are presented as mean ± SEM. **(B-E)** Statistical significance was assessed by paired t-test. ns, not significant; *, p < 0.05; **, p < 0.0005; ****, p < 0.0001. **(F)** Western blot analysis confirming efficient depletion of HECTD1 following siRNA treatment. Samples from three representative donors (#) are shown. β-actin was used as loading control.

**Figure S4 (related to Figure 4).**
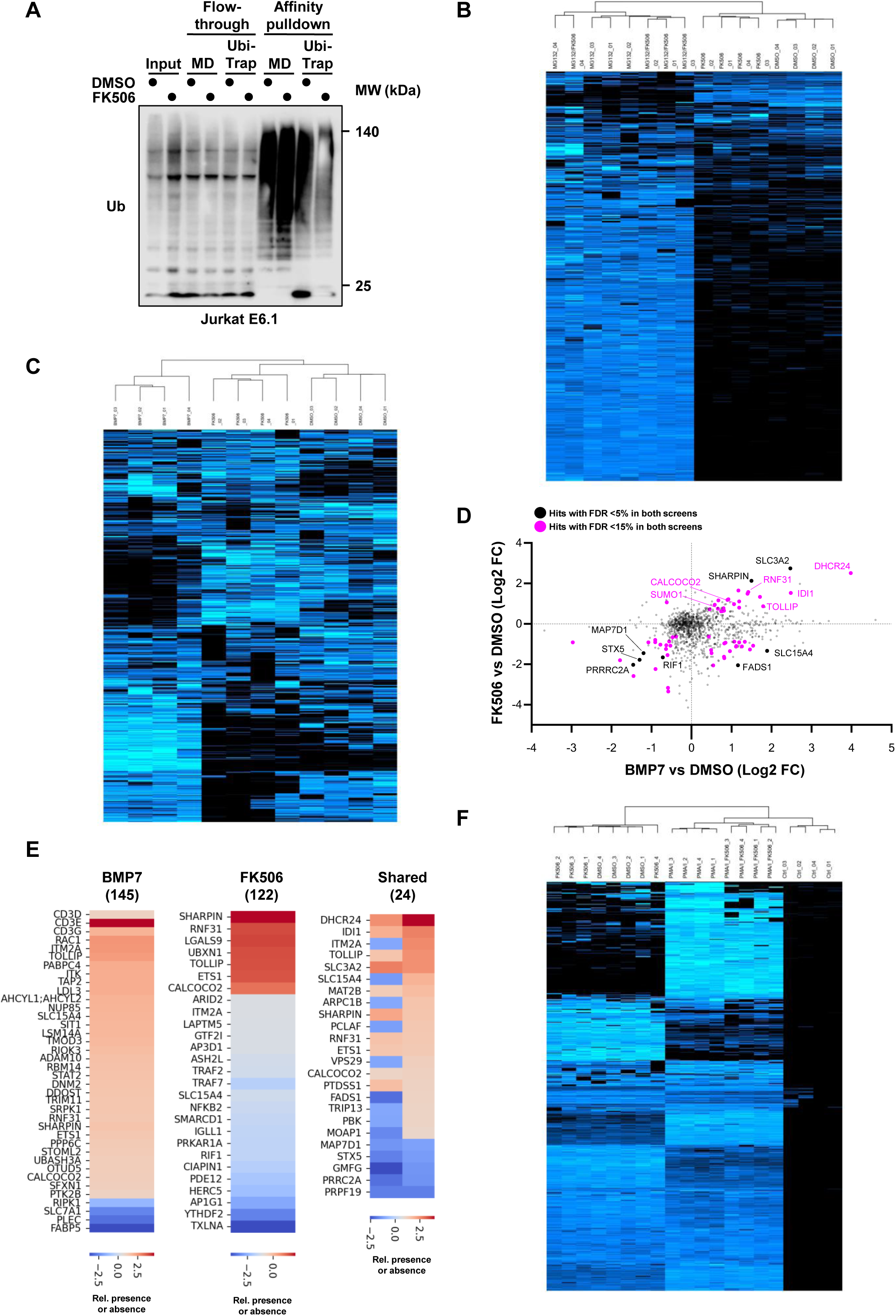
**(A)** Immunoblot analysis of ubiquitin affinity pulldown of protein extracts from Jurkat E6.1 cells treated with FK506 or mock-treated with DMSO, as indicated. MD, mock depletion. **(B-C)** Pearson-based unsupervised hierarchical clustering of z-scored ubiquitin affinity purification-mass spectrometry (AP-MS) datasets relative to Figures 4B (B) and 4C (C). Columns represent individual samples and rows represent detected proteins. Data were normalized by z-score prior to clustering. The heatmap shows relative abundance patterns across the samples, with black indicating low or no detectable signal and blue indicating increased relative abundance. Samples cluster according to similarity in protein association profiles, as indicated by the dendrogram on the right, highlighting condition-specific signatures. **(D) S**catter plot showing the relationship between ubiquitination changes induced by BMP7 and FK506 treatments relative to DMSO control, represented as log2 fold change (Log2 FC) values for each quantified ubiquitinated protein. Each dot corresponds to an individual ubiquitinated protein. Proteins significantly regulated in both screens are highlighted in black (FDR < 5%) or magenta (FDR < 15%). **(E)** Heatmaps showing selectively regulated ubiquitinated proteins identified following BMP7 or FK506 treatment relative to DMSO control. Numbers in parentheses indicate the total number of significantly regulated proteins in each treatment. Colour scale represents log2 fold change in ubiquitination abundance, with red indicating increased ubiquitination and blue indicating reduced ubiquitination relative to control. **(F)** Pearson-based unsupervised hierarchical clustering of z-scored ubiquitin AP-MS dataset related to Figure 4F. See panels B-C for a detailed description of heatmap clustering.

